# Machine learning behavioral analysis reveals cervical instability as an early biomarker of Amyotrophic Lateral Sclerosis

**DOI:** 10.1101/2025.09.29.679318

**Authors:** Omer Barkai, Xiangsunze Zeng, Keunjung Heo, Radhesh K Gupta, Natalie MacKinnon-Booth, Elke Bentley, Emily Shea, Clifford J Woolf

## Abstract

Early detection of neuromuscular disorders is a major clinical challenge, with most diagnoses only occurring after considerable motor neuron degeneration has already taken place. The central problem for early diagnosis of neuromuscular diseases is the subtlety of early symptoms and where to look for them. Without defined behavioral markers, the earliest stages of disease go undetected, delaying intervention and limiting neuroprotective therapeutic evaluation. Here, we present a machine learning (ML) based framework that identifies subtle postural alterations in freely behaving animals. Using longitudinal pose data from SOD1^G93A^ mice, a widely used Amyotrophic Lateral Sclerosis (ALS) mouse model, we focused on postural states in idle periods, behavioral states usually overlooked in disease monitoring. Our analyses revealed consistent deviations in posture and a feature analysis pinpointed cervical instability during adolescence as a key distinguishing feature. We validated these findings through two independent behavioral assays engaging cervical musculature: rearing and wet-dog shakes, both of which showed significant impairments in male SOD1^G93A^ mice as early as 3 weeks of age, many weeks earlier than conventional muscle function assays. This approach establishes an unbiased, non-invasive, scalable strategy for detecting early-stage neuromuscular dysfunction, and provides a foundation both for clinical behavioral biomarker development in ALS and related disorders and will enable evaluation of early neuroprotective interventions.

## Introduction

ALS is a progressive neurodegenerative disease marked by selective degeneration of upper and lower motor neurons, resulting in muscle weakness, paralysis, respiratory failure and early death^1,2^. The disease often begins with either limb weakness, known as ‘spinal onset’, or with difficulties in speech and swallowing, referred to as ‘bulbar onset’. However, by the time symptoms become clinically evident, motor neuron degeneration is often very advanced, limiting therapeutic options essentially to symptomatic relief rather than prevention of disease progression.

Although emerging interventions have shown some promise in delaying ALS progression there is a lack of screening methods to detect early motor-related signs of ALS to enable evaluation of early neuroprotective interventions^1,3,4^. While several molecular biomarkers and imaging techniques have been explored for early ALS detection, these methods remain either invasive, inconsistent, or inaccessible for routine screening ^5,6^. There is, therefore, a critical gap in examinations that can capture the earliest manifestations of disease. The mean patient survival from ALS symptom onset is 3-5 years, with currently available drugs allowing only a short increase in survival of 6-19 months^1^.

A well-studied preclinical model for studying ALS is the SOD1^G93A^ mouse, in which ALS-like symptoms are believed to only start in adulthood at approximately 9-13 weeks of age ^7–9^. These are identified using conventional muscle strength approaches like the rotarod test, quantitative strength and stability exams, and gait analysis. These impairments progress to pronounced muscle weakness and atrophy, ultimately culminating in paralysis and premature death at approximately 16–20 weeks^7–9^. Although mouse anatomical studies suggest the presence of early motor neuron defects in young individuals, no well-defined criteria exist for non-invasive diagnosis at these stages, a major diagnostic gap^10^. Identifying reliable, non-invasive markers capable of detecting disease in its pre-symptomatic/early phases is therefore a pressing and unmet need.

We present here an ML-derived framework for identifying early ALS behavior markers in the SOD1^G93A^ mouse model. Using pose-estimation-derived features and particular AI techniques we recorded freely behaving SOD1^G93A^ mice, continuously up to 16 weeks, focusing on idle periods for stable assessment of posture. We detected distinct postural signatures distinguishing SOD1^G93A^ mice from their littermate controls, specifically in males but not females, which resembles clinical ALS sex differences ^11,12^. SHAP feature analysis identified cervical positioning as the most informative discriminator, leading to a hypothesis that reduced neck strength represents an early marker of ALS symptoms^13^. Consistent with this prediction, male SOD1^G93A^ mice exhibited significant reductions in two neck-strength-sensitive assays, rearing and wet-dog shakes, as early as 3 weeks of age. This data-driven, unbiased framework uncovers, therefore, previously unrecognized identifiers of neuromuscular disease and holds translational potential for early diagnosis and temporal monitoring of neurodegenerative motor disorders.

## Results

### ML framework for identifying SOD1^G93A^ animals versus littermate controls based on idling body pose

To identify potential postural differences between SOD1^G93A^ mice and their littermate controls we used a bottom-up behavioral platform^14^. Mice were recorded during 30-minute free-behavior sessions from 2 to 16 weeks of age **(Figure 1A)**. Videos were analyzed frame-by-frame for active behavior using BAREfoot, a recently generated algorithm for automatic ML-based rodent behavior analysis^15^. To identify changes in body part positioning during idle periods, we analyzed changes in the animal’s pose when the animals were not engaged in active behavior or movement (>0.5 sec, see Methods).

**Figure 1.**
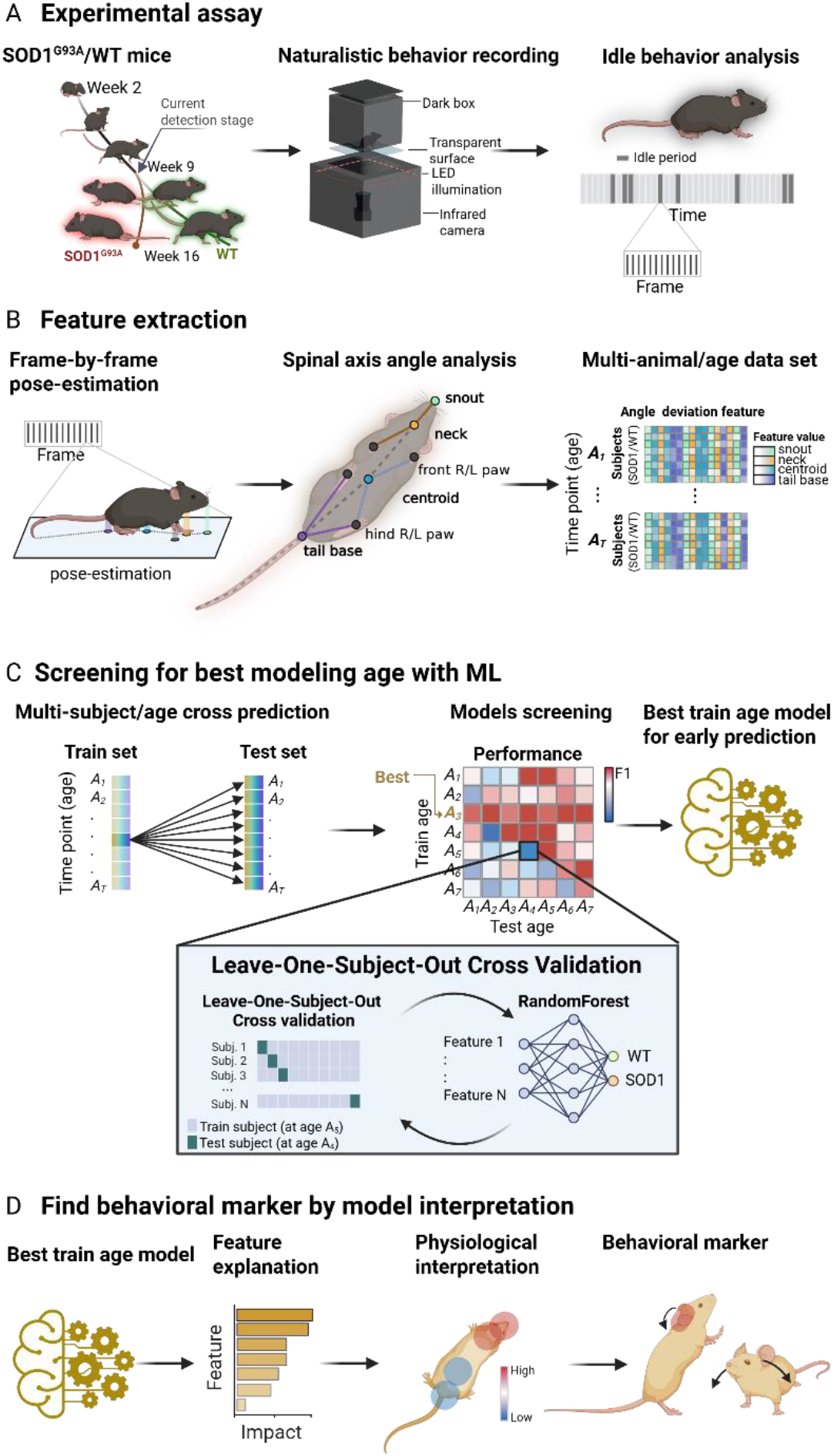
Workflow for pose-based SOD1^G93A^ animal classification. **(A)** Model of experimental assay. Mice from 2 to 16 weeks of age were recorded in a platform allowing them to freely move on a transparent surface, in the dark, while a NIR camera together with NIR (∼850nm) LED illumination captures body movements. Behavior is then analyzed post-hoc, and idle periods extracted accordingly. Mice with a red and green glow represent SOD1^G93A^ mice and wildtype mice at ALS-symptom development stage respectively. Mice without a glow represent the current limitation of detecting differences between SOD1^G93A^ and wildtype by behavior. The golden arrow indicates the currently available onset detection stage. **(B)** For every frame included in the idling period, a set of all the angles created between the spinal midline (snout, neck, centroid and tail base) and each of the key-point is extracted. Three example angles are shown for neck, centroid and tail base vertexes. Ray colors match vertex color. The standard deviation is then computed for every animal at each time point (age), creating a multi-animal/timepoint dataset. **(C)** The multi-animal/age data are split into train and test sets for a Leave-One-Subject-Out Cross Validation process. Training is done at every timepoint (Age: A_1_,A_2_,…A_T_) and tested at every timepoint (Age: A_1_,A_2_,…A_T_). An animal in the training set, regardless of the training timepoint, is always left out of the test set. The best age model for early prediction is then chosen for further analysis. **(D)** The best prediction model is analyzed with an explanatory feature analysis (using SHAP) to pinpoint the most informative body regions. According to the body features identified, relevant behaviors are then tested to validate their potential as an early behavioral biomarker.

Using DeepLabCut, we extracted animal pose dataset from every recording frame **(Figure 1B)**^16^. To capture posture changes we looked at the overall animal’s deviation in three-point body part angles, as such angular measurements are inherently scale-invariant and capture the relative orientation of body segments, which offers a robust basis for comparisons over time. To reduce feature dimensionality and isolate the most informative parameters for the model, we chose to include four midline-based body part angle vertexes (head, neck, centroid and tail base) that reflect movement symmetry with respect to the midline key points and the four limbs (left- and right-, front- and hind paws), and overall body orientation. Our primary data set included angles from the four vertexes, calculated for every video frame, each with seven complementary body parts, stacked and summarized to a per-video standard deviation of all included idle period frames (total of 84 features, 21 per 4 vertex categories). The final set of features include body part angle standard deviation which was created for every animal at every time point (age).

Early behavioral manifestations of ALS in SOD1^G93A^ mice are largely undefined. Instead, we focused on idle periods and performed a rigorous age-by-age statistical exploration to identify a high-performance model for distinguishing between SOD1^G93A^ mice and their wildtype littermates during these periods based on our feature set (**Figure 1C)**. To achieve this, we trained Random Forest-based ML models on age- and gender-based animal datasets. To ensure a robust and unbiased evaluation, a Leave-One-Subject-Out cross validation (LOSO-CV) approach was applied, such that no subject used for training was included in any test set. In LOSO-CV, the model is trained on all but one subject and tested on the excluded subject, repeating this process so that each subject serves as a test case once, providing a comprehensive assessment of generalizability across individuals. The selected features were used for training and testing the ML model over every timepoint.

After mapping all the trained models, the best performing model for early-stage classification can be identified and then analyzed with an explanatory feature analysis for interpretation of physiological translatability, pinpointing the body regions which impacted the model classification **(Figure 1D)**. These regions identify which early-stage behavioral tests could be used as behavioral biomarkers.

### Longitudinal ML Classification Reveals Early Disease Signatures in Male SOD1^G93A^ Mice

From birth to 4 weeks, mice are in an early postnatal phase, maternally dependent, rapidly developing motor and behavioral skills. Between 4 and 9 weeks, they enter adolescence, becoming independent, more aggressive, and sexually mature. By 9–10 weeks, mice reach adulthood with full behavioral and physical maturity **(Figure 2A)**^17^. Since the knowledge of symptoms evolving during pre-adulthood development stages are very limited, we used an exploratory approach by training Random Forest models for animals spanning from week 2 to week 16 of age and cross tested them on the same timepoints (ages), using LOSO-CV. To quantify the results for all the test sets, we computed the F1 scores comparing the targets to the ML predictions in males. A multi-animal/timepoint prediction heatmap was constructed to evaluate the fitted model’s overall performance in distinguishing SOD1^G93A^ and wildtype animals across ages **(Figure 2B, S1A)**. To verify that these results are not an outcome of chance, we repeated this process a 100 times with data shuffling the mice genotypes and comparing each shuffle result to the true unshuffled genotype’s result (see *Methods*). Overall, for males, models trained on adulthood stages (weeks 10–16) exhibit weaker generalization when tested at pre-adulthood stages, indicating that early disease stages may be characterized by distinct behavioral and physiological patterns. Adolescence-stage models, however, performed well for classifying animals at the adolescent test weeks. Notably, they also performed well in adulthood, suggesting that certain adolescent-stage features persist in adulthood.

**Figure 2.**
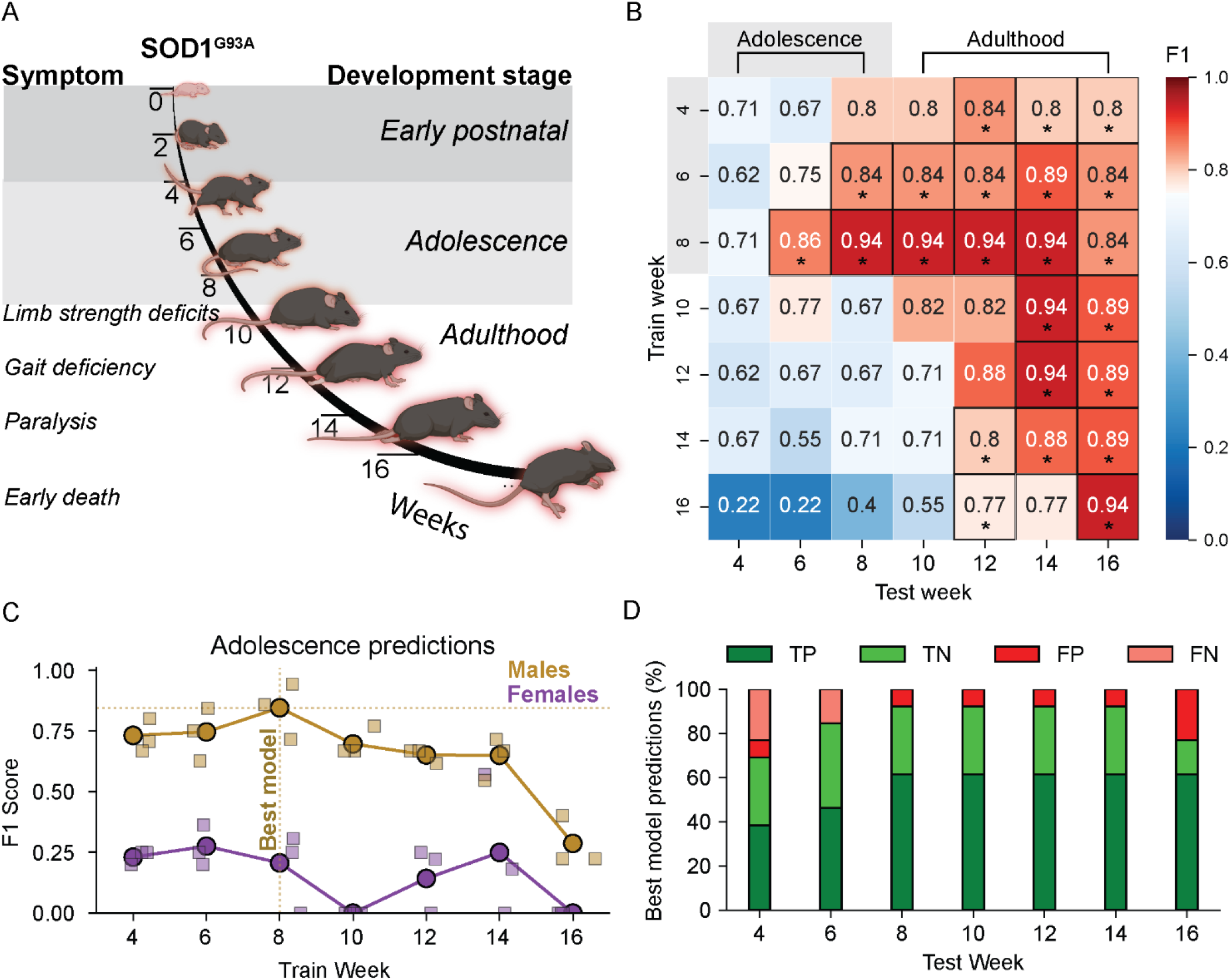
Multi-timepoint Leave-One-Subject-Out cross validation performance. **(A)** Illustration of ALS symptom development across mouse life stages with conventional detection methods. Grey shading depicts the time points for which non-invasive markers for ALS-like symptoms are not available. Shading: dark grey – early postnatal, light grey – adolescence, clear-adulthood. (**B**) heatmap showing the LOSO-CV performances (F1 score) of each train timepoint-to-test timepoint. Scores were compared to shuffled data. **(C)** Male (gold) and female (violet) mice adolescence (age 4-8 weeks) prediction results. Circles represent the F1 score for the merged test week data point predicted labels compared to the true labels. Squares represent the F1 scores from individual test weeks (4, 6, 8). The horizontal dashed line represents the best overall prediction (highest F1 score) and the vertical dashed line represents its corresponding training models timepoint. **(D)** Stacked bar graph showing the best cervical feature-based model’s prediction distribution by test weeks (TP – true positive, TN – true negative, FP – false positive, FN-false negative). * p <0 .05. N = 8 SOD1^G93A^ males, 5 wildtype males, 5 SOD1^G93A^ females, 9 wildtype females.

Next, we identified the optimal training timepoint for classifying SOD1^G93A^ vs. wildtype mice by evaluating the F1-score for the overall performance of each fitted model, comparing the stacked age prediction to the true genotypes (e.g. stacked week 4, 6, 8 predictions compared to their corresponding true labels) **(Figure 2C, S1B)**. Although fitted models trained on 2 to 3-week cohorts showed weak performance with low F1-scores, the male week 8 model achieved superior and consistent accuracy across multiple test weeks, suggesting that this timepoint captures critical disease-related features that generalize well both to adolescence stages and later stages **(Figure 2D; Figure S1A)**. The classification performance of week 8 models showed a first significant result for mice at the age of 6 weeks (F1-score: 0.86) and increased performance for the 8 weeks test set (F1-score: 0.94) **(Figure 2B, D)**. In sharp contrast to males, classification accuracy in female mice remained low across all ages **(Figure 2C; Figure S1B)**, suggesting that key early-stage behavioral and physiological changes emerge specifically only in male SOD1^G93A^ mice. Collectively, these findings reveal that male SOD1^G93A^ mice exhibit ALS-related behavioral changes far earlier than previously recognized, becoming detectable as early as 6 weeks of age.

**Supp. Figure 1.**
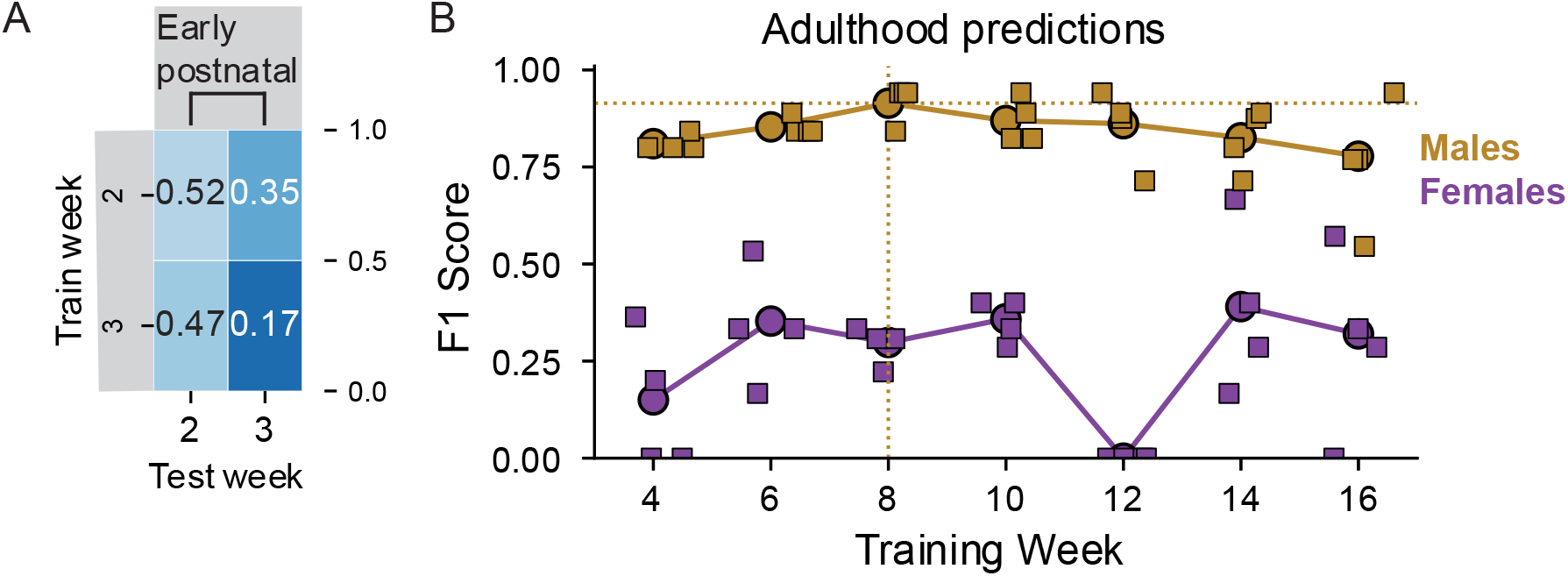
multi-timepoint Leave-One-Subject-Out Cross Validation performance for pre-wean and late-stage males, and early- and late-stage females. Related to Figure 2. (**A**) heatmap showing the LOSO-CV performances (F1 score) of each train timepoint-to-test timepoint for weeks 2-3. **(B)** Male (gold) and female (Violet) mice adulthood (age 10-16 weeks) prediction results. Circles represent the F1 score for the merged test week data point predicted labels compared to the true labels. Squares represent the F1 scores from individual test weeks (10,12,14,16). The horizontal dashed line represents the best overall prediction (highest F1 score) and the vertical dashed line represents its corresponding training models timepoint. N = 14 SOD1^G93A^, 14 wildtypes.

### ML feature importance analysis identifies the cervical region as an early-stage ALS-related behavioral locus

To gain physiological insight into early signs of ALS, we examined which dataset features were most influential in driving a high ML classification performance. To this end we used the SHAP (Shapley additive explanations) analysis toolkit, an explainable AI technique for analyzing feature contributions to the model’s decision, to determine the features of the best-performing model, based on its predictions^13^. For each LOSO training session, we utilized the SHAP explainer to analyze feature contributions and importance for the performance over the iteration. The SHAP values for the best model, trained with week 8’s data (see Figure 2C), were then combined, providing an explanatory ranking of feature importance, which we then categorized to the four groups of vertexes along the body axis: snout, neck, centroid and tail base. Importance ranking showed the highest values for cervical positioning (snout and neck) related features **(Figure 3A, Figure S2A)**. SHAP analysis indicates that for the top importance features of the snout and neck, larger deviations in angles are associated with wildtype animals, indicating more cervical-related vertex flexibility. In an overall feature-by-feature SHAP-based ranking, 50 percent of the feature weights were assigned to the snout and neck **(Figure S2A)**. We further explored the importance of key spinal midline categories (feature vertexes) and their connection to the limb nodes, by summing the importance of each key point importance weights and connection weight, respectively. Overall, the snout and neck (cervical region) features had more impact compared to the centroid and tail base (lumbar region) features **(Figure 3B)**. The cervical region’s connection to the front limbs and lumbar region were stronger than to the hind limbs.

**Figure 3.**
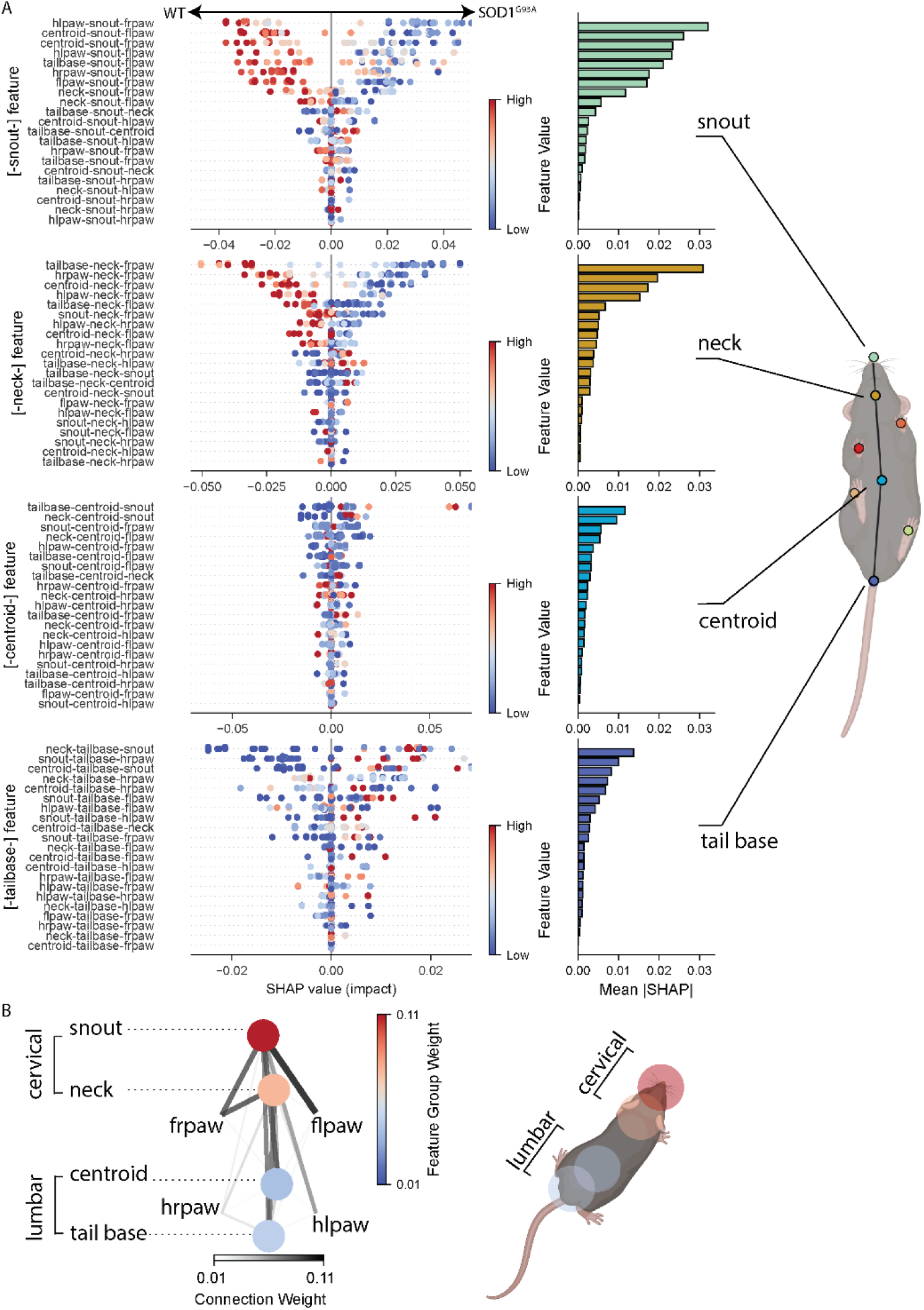
Feature importance identification in SOD1^G93A^ versus littermate control mice. (A) SHAP feature importance categorized by midline vertexes, ranked from the highest to lowest importance by category. Left, SHAP plot illustrating how each feature influences the prediction for SOD1^G93A^ (right-shifted) or wildtype (left-shifted). Each point represents a test sample’s SHAP value for a specific feature, indicating the feature’s contribution to the model’s output. A point represents the contribution of a feature per LOSO iteration. Thirty randomly chosen points were stacked vertically to reflect the density of SHAP values per feature. The color of each dot corresponds to the original feature value—blue for low and red for high—scaled according to that feature’s range in the dataset. For example, if blue points cluster on the right side of the plot for a given feature, it suggests that lower values of that feature are associated with an increased likelihood of being a SOD1^G93A^ animal. Right, the overall importance of each feature by category, descending from top to bottom. Value achieved by averaging the absolute SHAP values of each feature. (B) Left, summation map of the model midline vertex feature weights, and their limb connection weights. Right, the midline vertex summary map are shown on a mouse illustration.

**Supp. Figure 2.**
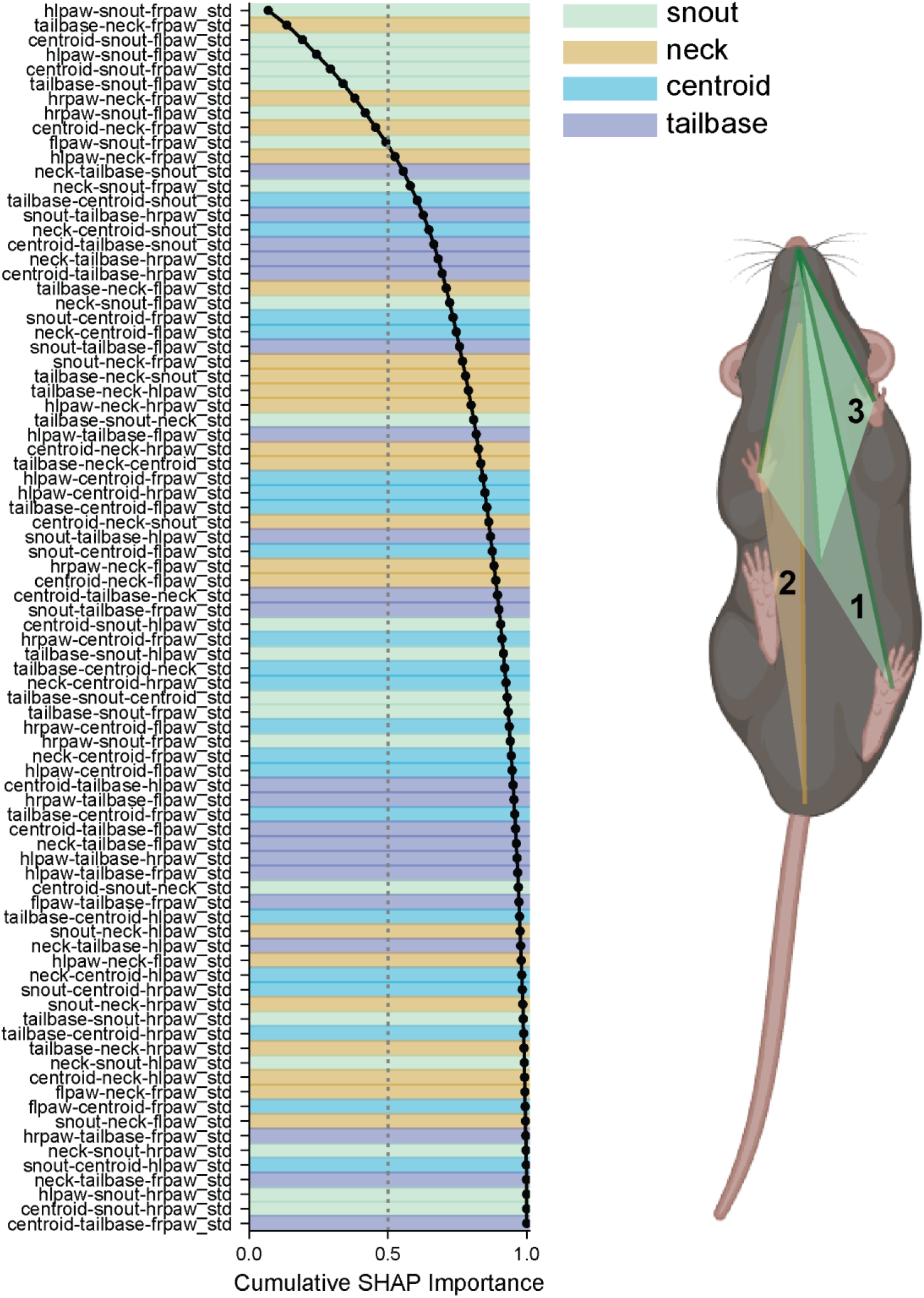
Relative feature weight by category. Related to Figure 3. Left, feature weights normalized to the sum and sorted by relative weight descending from highest (top) to lowest (bottom). The black curve represents the cumulative sum added by every normalized feature weight. Background colors represent the feature category. Right, the first 3 features displayed on of a bottom-up mouse illustration. The numbers on the angles indicate the feature’s importance rank.

### Cervical features predict SOD1^G93A^ mice at adolescence stage

Our feature analysis shows that the cervical region has a high impact on the model’s performance to discriminate SOD1^G93A^ mutants. This led us to hypothesize that the cervical region could serve as a strong candidate for early-stage diagnosis of the disease. We first tested if cervical features (snout and neck) alone are sufficient for early prediction of ALS-symptoms in SOD1^G93A^ mice by excluding the lumbar features (centroid and tail base) (**Figure 4A**). For comparison, we also tested the lumbar feature-only model in the absence of cervical features **(Figure 4B, C)**. Only a single train/test set performance for early prediction was significant compared to the shuffled targets results (**Figure 4B**). The best cervical feature-based model, as in the full model, was also found when trained on week 8 data with a first significant peak testing on 8-week data (F1 score: 94) **(Figure 4A, C, D; Figure S3A)**. These data strengthen our conclusions that the cervical region is key for identifying early-stage impairments in SOD1^G93A^ mice.

**Figure 4.**
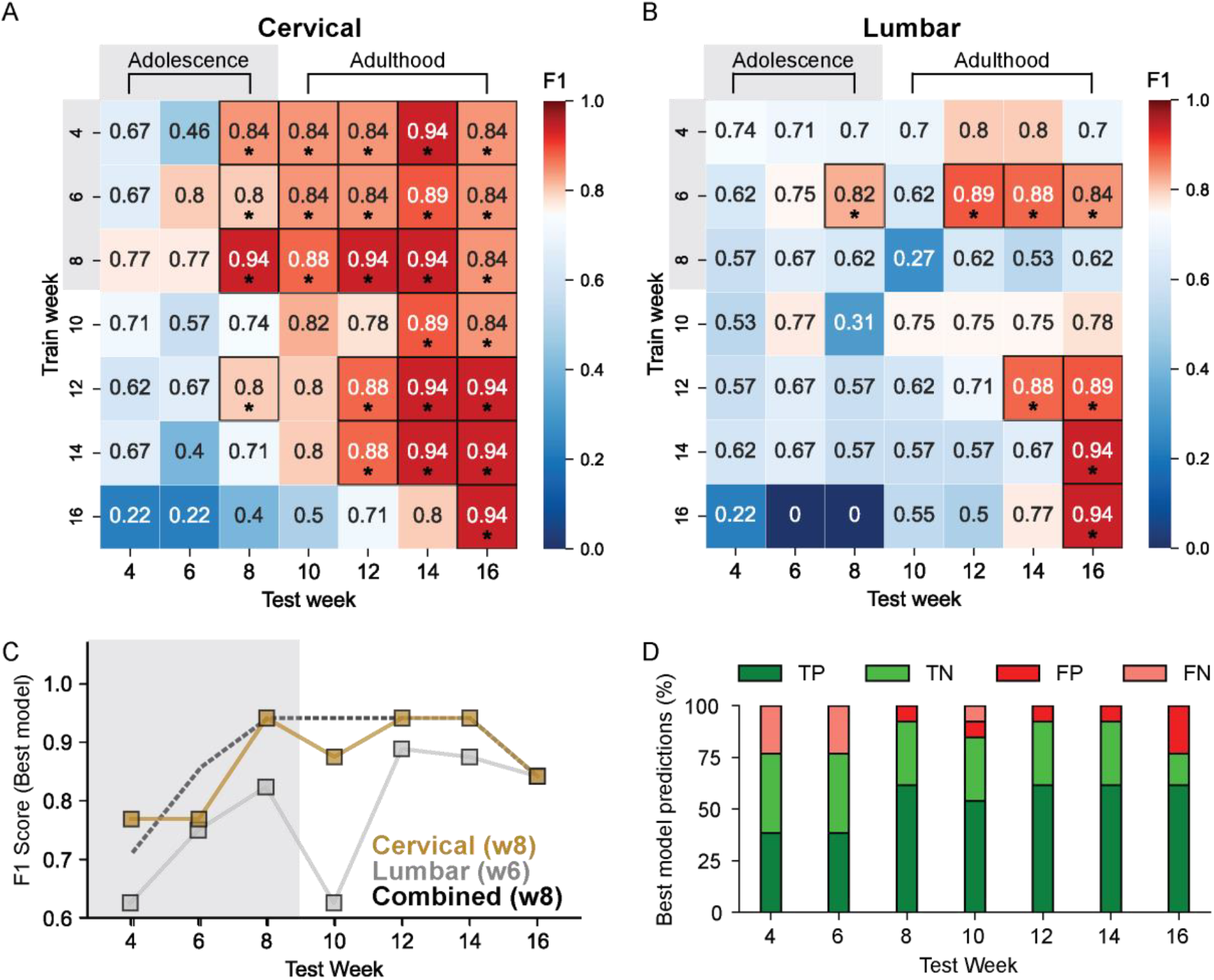
Cervical-versus lumbar-only features for SOD1^G93A^ prediction. **(A)** Heatmap showing the LOSO-CV performances (F1 score) of each train timepoint-to-test timepoint with Cervical-only features. **(B)** As in A, with Lumbar-only features. Only a single train/test set performance for early prediction was significant when compared to the shuffled targets results with overall two above 0.75 F1-score, compared to four in cervical and seven above 0.75 F1-score. **(C)** Cervical, lumbar and combination (adapted from Figure 2) best models (train weeks 8, 6 and 8, respectively) prediction results. Gold, silver and black represent cervical, lumbar and combined respectively. Squares represent the F1 scores achieved by the best models found in C. Light grey shading depicts adolescence stage. The horizontal dashed line represents the best overall prediction (highest F1 score) and the vertical dashed line represents its corresponding training models timepoint. **(D)** Stacked bar graph showing the best cervical feature-based model’s prediction distribution by test weeks (TP – true positive, TN – true negative, FP – false positive, FN-false negative).

**Supp Figure 3.**
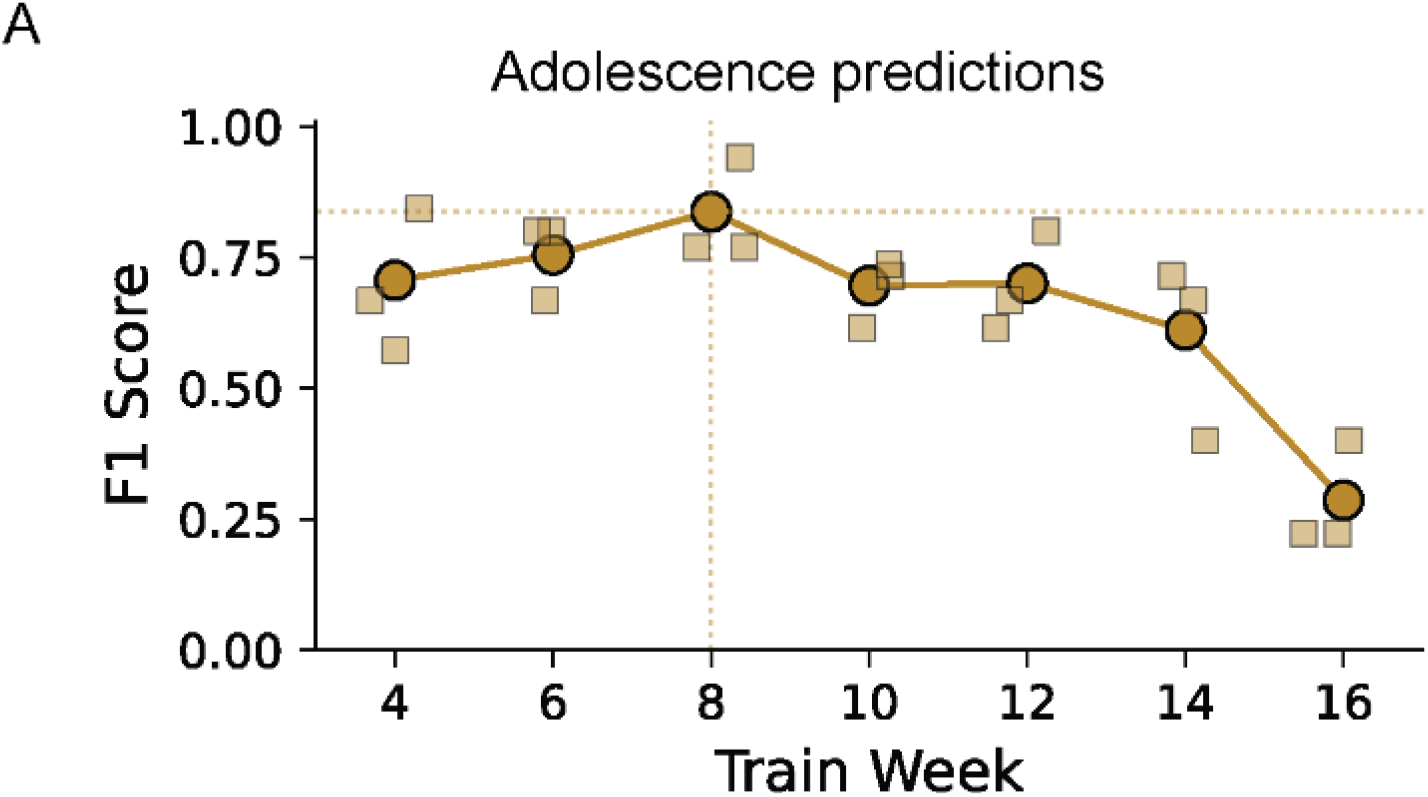
SOD1^G93A^ prediction performance during adolescence with cervical features. Related to Figure 4. (A) Adolescence (age 4-8 weeks) prediction results summary for cervical-only feature-based predictions. Circles represent the F1 score for the merged test weeks data point predicted labels compared to the true labels. Squares represent individual test week (4,6,8) time point’s F1. The horizontal dashed line represents the best overall prediction (highest F1 score), and the vertical dashed line represents its corresponding training models timepoint, colored with respect to the plot.

### Cervical-Related Motor Behaviors Are Impaired from Early Stages in SOD1^G93A^ mice

Neck muscle weakness is a well-recognized symptom in late ALS-development in humans, often leading to a head drop^18–23^. We reasoned that changes in cervical movement may also be evident in early-stage physiological assessment. To test this, we performed a behavioral analysis using the BAREfoot pipeline. We first measured rearing, an exploratory behavior that strongly engages cervical muscles as the animal raises its head while upright on its hind limbs **(Figure 5A)**. SOD1^G93A^ mice engaged in less rearing behavior compared to their wildtype littermates throughout the full adolescence period with age-by-age analysis and per-subject cumulative summation examinations **(Figure 5B, C)**. Because rearing requires hindlimb weight bearing, reduced rearing time could potentially reflect hindlimb weakness rather than cervical deficits. To address this, we examined grooming bouts, during which mice also support weight on the lumbar region while nibbling, licking, and rubbing the body and face. However, grooming behavior did not differ significantly between SOD1^G93A^ and wildtype mice across timepoints, supporting the conclusion that reduced rearing is attributable to a cervical rather than a hindlimb weakness **(Figure S4A-B)**.

**Figure 5.**
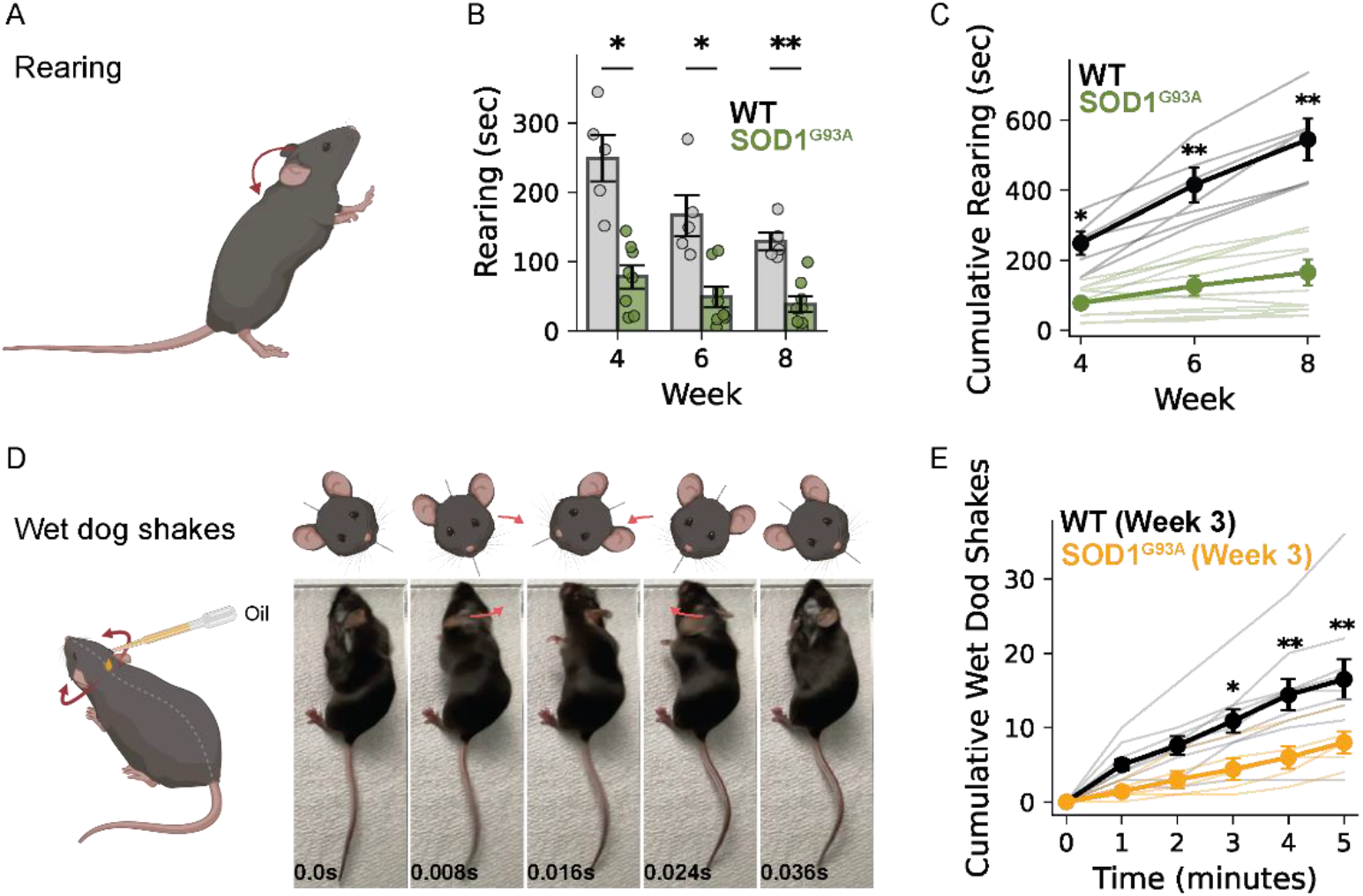
Impaired neck-related activity in SOD1^G93A^ animals. **(A)** Illustration of mouse rearing behavior. The red arrow represents the head lifting direction. **(B)** Quantification of rearing behavior for early-stage time points recordings (total time per 25 min). **(C)** Cumulative summation of B. N = 8 SOD1^G93A^, 5 wildtypes. **(D)** Left, illustration of the oil-drop experimental setup for evoking wet dog shakes. Right, a representative cycle (left-right-left) from a four-cycle wet dog shake bout. The illustrations show head turn direction. Red arrows show the head turning direction in the frame. (**E**) Cumulative number of wet dog shake bouts in wildtype vs. SOD1^G93A^ animals recorded from 3-week-old mice. * p <0 .05, ** p<0.01. N = 5 SOD1^G93A^ males, 10 wildtype males. Shown are the Mean ± SEM. Dots and bright shaded lines refer to individual animals

While a significant difference in rearing time was found at 4 weeks of age, a trend was observed even earlier, during early postnatal stages, at weeks 2-3 **(Figure S4C-D)**. We therefore sought to further investigate behavioral changes during the early postnatal period by using a more precise and spatiotemporal controlled cervical strength test. Previous work showed that sunflower seed oil applied to the neck elicits repeated bouts of wet dog shakes, characterized by rapid and strong back-and-forth oscillatory turn (∼3 times at ∼19 Hz) of the cervical region **(Figure 5D)**^24^. We applied a droplet of oil to the dorsal neck of 3-week-old mice. While this stimulus reliably triggered repeated wet-dog shake bouts in all animals, SOD1^G93A^ mutants exhibited significantly fewer events than their wildtype littermates over the 5-minute recording period **(Figure 5E)**.

Taken together, these analyses identify the cervical region as a critical driver of early identification of SOD1^G93A^ mice and demonstrate that cervical-related behaviors are impaired in these mice from as early as 3 weeks of age, well preceding the current known adulthood muscle weakness symptom onset markers which only manifest at 9 or more weeks.

**Supp Figure 4.**
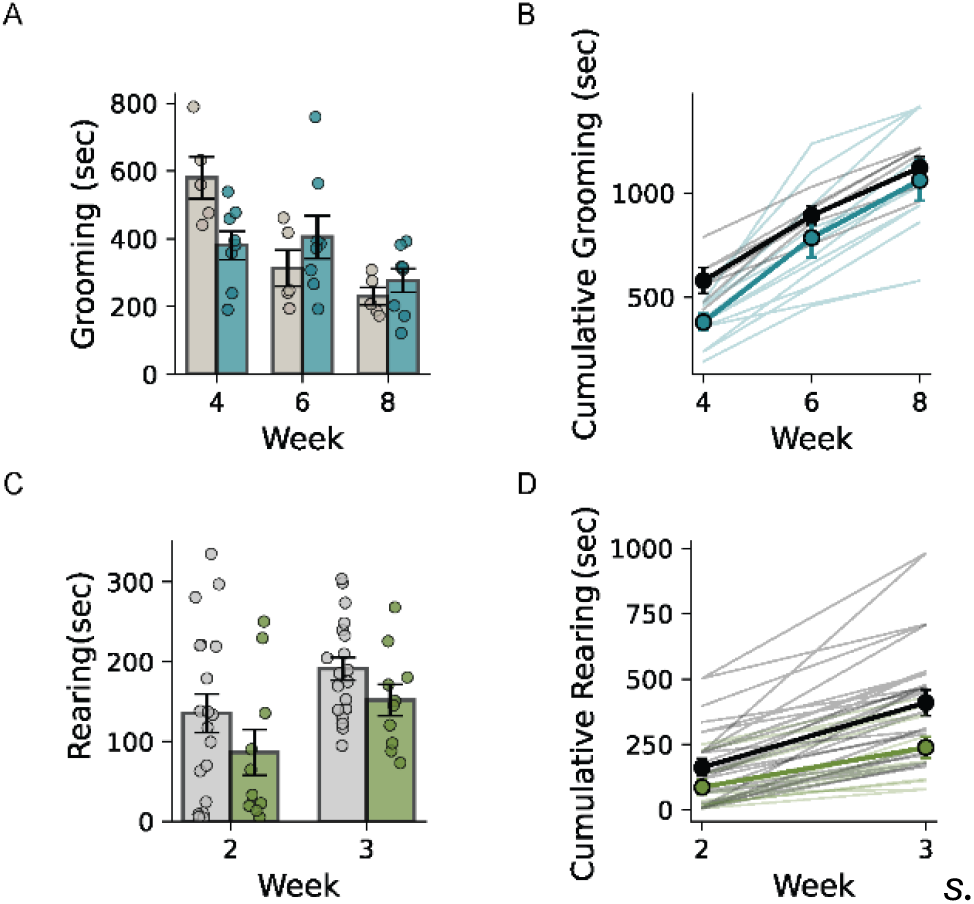
Adolescence grooming and early postnatal rearing in SOD1^G93A^ mice. Related to Figure 5. (A) Quantification of grooming behavior for adolescence stage time points recordings (total time per 25 min).(B) Cumulative summation of B. N = 8 SOD1^G93A^, 5 wildtypes. (C) Quantification of rearing behavior for early postnatal stage time points recordings (total time per 25 min). (D) Cumulative summation of D. N = 10 SOD1^G93A^, 19 wildtypes. Shown are the Mean ± SEM. Dots and bright shaded lines refer to individual animals.

## Discussion

In this study, using pose-estimation-derived features and AI techniques, we identified the cervical region, an often-overlooked site of disease onset, as highly informative for early detection of disease in SOD1^G93A^ mice. We present an ML-based approach for early-stage identification of ALS-like phenotypes that does not rely on predefined behaviors or anatomical assumptions. Instead, our data-driven framework enables unbiased screening for physiological signatures. By decomposing the most informative features, we generated hypotheses about early behavioral changes and, thereby, shifted the window for detecting motor decline from adulthood to early postnatal and adolescent stages. Follow-up behavioral assays that primarily engage neck muscles validated this result, reinforcing the translational potential of these findings.

Our pipeline allows for the discovery of candidate regions without relying on preconceived assumptions. This is in contrast to most prior studies of ALS that begin with a focus on known clinical features, such as limb weakness or gait abnormalities, which typically emerge later, during adulthood disease stages. Our SHAP-based analysis revealed that small but consistent deviations in cervical posture, imperceptible to the human eye, can serve as early indicators of motor dysfunction. This highlights the power of explainable AI not only for prediction, but also for directing disease mechanism investigations in a hypothesis-generating fashion.

Previous literature on the SOD1^G93A^ mouse model has described symptom onset only at adulthood, typically 9–13 weeks of age, with detectable changes in rotarod performance, grip strength, and gait occurring at or after this period. Here, we demonstrate that spontaneous behavioral differences between SOD1^G93A^ and wildtype animals are already present during adolescence, onsetting at 4 weeks when assessed with rearing. We further show behavioral differences at the early postnatal stage, as early as 3 weeks when assessed through evoked wet dog shakes. This significantly advances the temporal window for disease detection, opening new possibilities for interventional and mechanistic studies in the earliest phases of disease.

ALS onset is commonly associated with weakness in the limbs, or speech and swallowing difficulties^2,25^. Neck muscle weakness is also observed in advanced stages of disease, often manifesting as head drop due to cervical extensor weakness^18–23^. Our mouse findings suggest that analogous muscles in humans may also be functionally compromised early in disease development. While neck muscle strength is rarely assessed in early-stage human ALS, our results support exploring such measurements through targeted strength testing, posture analysis or wearable motion sensors, into potential diagnostic or monitoring frameworks.

Interestingly, the observed predictive features and behavioral differences were specific to male SOD1 ^G93A^ mice. This aligns with prior studies reporting more robust phenotypes and accelerated progression in male animals, potentially mirroring the higher incidence of ALS in human males^2,8,9^. However, the absence of similar findings in female mice also highlights sex-specific disease mechanisms and model generalizability. Further investigation into sex differences, both in mice and in human ALS, will be essential for developing inclusive diagnostic strategies.

This study presents a novel unbiased framework, and rigorous approach for uncovering early behavioral signatures across models of neuromuscular and neurodegenerative disorders. Our findings specifically highlight the diagnostic value of impaired cervical motor control in ALS and demonstrate the broad utility of explainable AI as a tool for hypothesis generation in biomedical research. Future work can now determine whether interventions applied during this newly defined pre-symptomatic window can slow or prevent disease progression, thereby offering a neuroprotective strategy rather than merely halting advanced disease progression or alleviating established symptoms.

## Methods

### Animals

All experiments were performed in a blinded fashion and in strict accordance with the guidelines of the Boston Children’s Hospital Institutional Animal Care and Use Committee (protocol no. 00002403). Hemizygous SOD1^G93A^ mice (B6SJL-Tg(SOD1*G93A)1Gur/J) were obtained from The Jackson Laboratory (JAX#002726), and maintained by breeding hemizygous males with C57BL/6 wildtype females (JAX#000664). This breeding scheme generated both transgenic heterozygotes and wildtype littermates, which were distinguished by PCR genotyping using primers specific to the human SOD1^G93A^ transgene, and further confirmed by the presence of hindlimb paralysis at the endpoint (16 weeks of age). All mice were housed under standard conditions on a 12-h light/dark cycle at 21–23 °C with 30–50% humidity.

### Behavioral platform and video recording

The recording setup, using NIR light together with frustrated total internal reflection technology (FTIR), was described in detail by Zhang et al.^14^, however, in this study FTIR was not used. Image brightness features were extracted using image analysis tools utilizing LED light-reflected changes in pixel intensity. Briefly, an 18 × 18 × 15 cm (length x width x height) black acrylic box closed on all sides except for the bottom was placed on a 25 cm square piece of 5-mm thick glass floor. A black acrylic plate was installed under the camera, to prevent entrance of light below the glass and a 25 × 25 cm black acrylic surround frame panel with an 18 × 18 cm square cutout was positioned on top of the glass floor panel to prevent light leaking to the camera from above (**Figure 1A**). Two separate 850-nm NIR LED strips (SMD5050-300-IR, Huake Light Electronics Co, Ltd, Shenzhen, China) were positioned horizontally 10 cm below the glass floor to provide illumination of the animals from below and positioned such that reflections from the LEDs off the sides, top, or floor of the chamber were not visible from the camera position. Power to all LEDs was provided by a 12-V DC power supply. To record the animals in the dark an NIR camera (Basler acA2000 -50gmNIR GigE, Basler AG, Ahrensburg, Germany) was positioned 30 cm beneath the glass surface. Pylon viewer software (v4.1.0.3660 Basler) was used for the video recordings. The same camera settings parameters were used for each recording. Videos were acquired at 25 Hz with 1000 × 1000 pixels dimensions and then downsized to 512 × 512 pixels for analysis using ImageJ’s scaling function, specifying to average during the downsizing.

### Body part tracking

Automatic body part tracking (pose estimation) was done as described in Zhang et al. ^14^ and Barkai et al.^26^. Briefly, video frames from recordings of mice in the behavioral platform were randomly selected. Nine key body parts (four paws, snout, neck, centroid, tail base and tail end) were manually labeled using DLC. These were utilized to train a DLC model for mouse pose estimation. To address model errors caused by background interferences such as urine, feces, or light reflection, the training data included videos with these potential disruptions in the field of view. The trained model was applied to automatically locate all nine body parts of interest in a video. In this study, the tail tip was excluded as its length and high degree of freedom, making it highly variable in position and movement with respect to the animal’s posture. Furthermore, humans do not share a functional analog mice tail.

### Multi-animal/timepoint

#### Behavioral Feature Extraction from Freely Moving Mice

Videos of individually tracked animals from multiple cohorts spanning birth to sixteen weeks of age were recorded. A single video of 30 minutes, at 25 frames per second, was collected for each animal at every age timepoint.

Pose estimation data were extracted to .h5 files using DeepLabCut. The DeepLabCut estimations were filtered with a median filter type with a window of 7 frames. data were imported into Python and processed with a custom script as part of the feature extraction process. For every frame, we derived a set of geometric features that included the angles between every set of three key body parts, in which the spinal midline key points (snout, neck, centroid, tail base) are their vertex.

Only idle periods were retained for downstream analysis. To ensure that the animal was behaviorally idle in a frame, the maximum two-frame-difference velocity (*‘[body part]_vel2’*) summation across all body key points in that frame remained below a set threshold (≤2), and persisted for at least 0.5 seconds (i.e., 13 consecutive frames). finalfeature set per animal, at each age timepoint, was the calculation of standard deviation of all the key point body angles during idling behavior frames.

#### Classification Framework and Leave-One-Subject-Out Cross-Validation

To evaluate whether early behavioral signatures could distinguish SOD1^G93A^ mice from wildtype littermates, we implemented a Leave-One-Subject-Out cross validation (LOSO-CV) framework across all combinations of training and testing time points.

For each training time point (age) *t*_*i*_ (*i* = *A*_1_, *A*_2_, … *A*_*T*_), behavioral features from all animals of that age were pooled across cohorts. A Random Forest classifier (n_estimators=100, max_depth=2, class_weight=‘balanced’, random_state=42) was trained using LOSO-CV. Classifier threshold was set to the default (0.5). In each fold, all animals except one were used for training, and the held-out animal served as the test subject. This procedure was repeated until each animal had served exactly once as the test subject. Importantly, the held-out animal was excluded from training across *all* time points, preventing information leakage from overlapping behavioral patterns at different ages.

The trained model from each LOSO fold at *t*_*i*_ was then used to predict the genotype of the held-out animal at every possible testing time point *t*_*j*_ (*j* = *A*_1_, *A*_2_, … *A*_*T*_). Binary predictions were obtained from genotype probability scores using a threshold of 0.5.

For each training–testing pair (*t*_*i*_ → *t*_*j*_), classification performance was quantified by the F1-score, averaged across all LOSO folds. This process yielded an *A*_*T*_ × *A*_*T*_ performance matrix (see Figures 2B, 4A–B), where entry 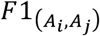 reflects how well a model trained at age *A*_*i*_ generalizes to data from age *A*_*j*_ . This design captures both within-age classification performance and cross-age generalization, enabling assessment of when disease-related behavioral signatures first emerge.

F1-score was measured as follows:

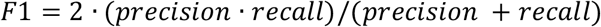

Where

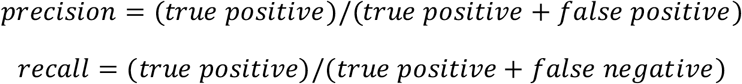

#### Statistical Evaluation of Classifier Performance

To assess whether the model’s performance (F1 score) was significantly better than chance, we conducted a permutation test. Specifically, we permuted the true class labels (*y*) 100 times, retrained the model on each permuted dataset, and evaluated the F1 score on the same input features (*X*). This procedure generated a null distribution of F1 scores representing performance expected under the null hypothesis of no relationship between features and labels.

The p-value was then computed as the proportion of shuffled F1 scores that were equal to or greater than the F1 score obtained using the original (non-permuted) labels. The final *p-value* was calculated as:

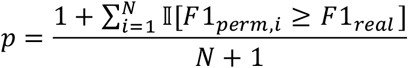

where *N* is the number of permutations (here, 100), *F*1_*real*_ is the F1 score from the model trained on true labels, *F*1_*prem,i*_ is the F1 score from the *i*^*th*^ permutation, and 𝕀 is the indicator function (1 if the condition is true, 0 otherwise).

#### SHAP-Based Feature Importance Analysis

To identify the most informative behavioral features associated with SOD1^G93A^ classification, we computed SHAP (SHapley Additive exPlanations) values for each LOSO training set using TreeExplainer for Random Forests ^13^. For each LOSO iteration, SHAP values from all folds were aggregated, and mean absolute SHAP values were computed across all animals and features. This yielded an age-resolved map of feature importance, revealing which features contributed most to classification at each stage of disease progression.

#### Rearing behavioral scoring

An automatic classifier for rearing behavior was built using the BAREfoot pipeline^26^. Thirteen videos of freely behaving mice, of different sexes and different ages, in a dark bottom-up behavioral platform were manually annotated for rearing behavior. A classifier was then tuned, trained and evaluated in a 5-fold cross validation and verified on a randomly chosen test set for performance levels. The final performance reached a high F1 score (≥ 0.9). After the classifier was built, all videos included in the study went through frame-by-frame rearing behavior automatic classification and analysis.

#### Wet dog shakes scoring

Wet-dog shakes were assessed as previously described (PMID: 39509513), with minor modifications. Briefly, animals were placed in a transparent acrylic chamber (20 × 20 × 15 cm; length × width × height) and recorded from above using a high-speed camera at 120 Hz. A 20-µl droplet of sunflower seed oil (Sigma, #S5007) was applied to the dorsal neck region with a pipette, after which mice were immediately returned to the chamber and recorded for 5 minutes. Wet-dog shake events were manually annotated and quantified from the 120-fps video recordings.

#### Data analysis computer specifications

All analyses were performed in Python 3.10 using custom code along with standard scientific computing libraries. Feature extraction and preprocessing were performed using BAREfoot’s feature extraction function.

Automated behavior scoring was performed with BAREfoot, implemented in Python (v3.10), in PyCharm (v2022.3.2) integrated development environment (JetBrains r.s.o, Prague, Czechia).

All analyses were done using a consumer-grade PC with an AMD Ryzen Threadripper 3970 x 32-Core Processor, 3700 Mhz, 128 GB RAM, and NVIDIA GeForce RTX3060 GPU, run on Microsoft Windows 10 operating system.

#### Statistics

Unless otherwise stated, statistical analysis was performed using two-way analysis of variance followed by Bonferroni correction for pairwise comparisons.

## Acknowledgements

This project was funded by the SMA Foundation (CJW).

